# Conservation and structural analysis of the Xenopus laevis phospho-proteome

**DOI:** 10.1101/009100

**Authors:** Jeffrey R. Johnson, Silvia D. Santos, Tasha Johnson, Ursula Pieper, Andrej Sali, Nevan J. Krogan, Pedro Beltrao

## Abstract

The African clawed frog *Xenopus laevis* is an important model organism for studies in developmental and cell biology, including cell-signaling. However, our knowledge of *X. laevis* protein post-translational modifications remains scarce. Here, we used a mass spectrometry-based approach to survey the phosphoproteome of this species, compiling a list of 3225 phosphosites. We used this resource to study the conservation between the phosphoproteomes of *X. laevis* and 13 other species. We found that the degree of conservation of phosphorylation across species is predictive of sites with known molecular function, kinase interactions and functionally relevant phospho-regulatory interactions. In addition, using comparative protein structure models, we find that phosphosites within structured domains tend to be located at positions with high conformational flexibility. A fraction of sites appear to occur in inaccessible positions and have the potential to regulate protein conformation.

## Introduction

Protein function can be regulated by post-translational modifications (PTMs) by altering diverse protein properties such as their localization, activity or interactions. Protein phosphorylation is one of the most well studied PTMs with over 50 years of research since the pioneering work of Krebs and Fischer on glycogen phosphorylase [1]. It is estimated that approximately 30% of the human proteome can be phosphorylated and this modification has been shown to play a role in a very broad set of cellular and developmental functions as well as dysregulation in disease [2]. Recent advances in phosphoenrichment procedures and mass spectrometry (MS) technologies have resulted in a tremendous increase in the capacity to identify phosphorylation sites on a large scale [3] and over the past few years over 200.000 phosphorylation sites have been identified across a varied number of species (ptmfunc.com). These studies have highlighted the true extent and complexity of PTM regulation and underscore the need to develop large-scale approaches to study PTM function. Making use of these phosphorylation, data evolutionary studies have revealed that phosphosites tend to be poorly constrained and a large fraction are not conserved across species [4–9]. Given the high evolutionary turn-over of these modification sites it is plausible that a fraction of these serve no biological purpose in extant species [4,10]. Therefore, it has become important to develop methods to discern the functional relevance of PTMs [11]. For example, the conservation of phosphosites has been used to highlight sites that are more likely to be important. Different kinases have specific preferences for the amino acids in the vicinity of the target phosphorylated residue [12–14]. This local sequence context is often referred to as the kinase target consensus sequence or motif and the conservation of these kinase motifs across orthologous proteins can be used to improve the predictions of kinase regulated sites [15,16]. In parallel to conservation based approaches, computational and experimental methods have been developed to identify phosphosites that are more likely to be functionally important by regulating protein interactions [10,17], protein activities [10], metabolic enzymes [18], or cross-regulate other types of modifications [19–21].

Phosphoproteomic approaches have been applied extensively to several species [6,7,22–25]. However, although *X. laevis* is a well-established model organism there has been little previous knowledge of the extent and conservation of its phosphoproteome. To address this we have used a MS approach identify phosphorylation sites in *X. laevis* egg extracts. This approach resulted in the identification of 2852 phosphorylation sites. For the subsequent analysis, we combined these sites with sites identified in a recent study [26] resulting in a total of 3225 phosphosites for analysis. Using a compilation of phosphorylation information for 13 other species we identified a small number of highly conserved phosphosites, which were found to be enriched in sites with known molecular functions. In addition we used kinase specificity predictions for kinases involved in cell cycle regulation to predict conserved kinase-protein associations. The degree of conservation of predicted cell cycle-related kinase interactions was correlated with known kinase-protein regulatory interactions. Conserved putative interactions were also enriched in proteins that are phosphoregulated during the cell cycle and in genes that when knocked down cause mitotic phenotypes. In order to study the structural properties of these sites we obtained comparative models for 1239 phosphoproteins. Structural analysis revealed a number of solvent inaccessible phosphosites that likely indicate protein regions that can exist in more accessible conformations. The analysis of these sites suggests that a significant fraction may regulate protein conformation. Beyond the structural and evolutionary observations, the phosphosites determined here provide a resource for future signaling studies in *X. laevis*.

## Results

### Conservation and structural coverage of the *X. laevis* phosphoproteome

Xenopus laevis egg extracts were prepared in two cell cycle stages (interphase and mitosis) in order to increase the coverage of phospho-regulatory events. After protein extraction the samples were trypsin digested, subjected to a phosphopeptide enrichment protocol and LC-MS analysis (see Materials and Methods). The spectra were matched to a reference proteome for *X. laevis* and a total of 820 and 1900 non-redundant phosphosites were obtained for the interphase and for the mitotic sample respectively. In addition we have also compiled 481 sites from a previous study [26] that we were able to map to the same reference proteome. The two sets obtained in this study have a total of 2852 non-redundant sites and the 3 combined sets have 3225 non-redundant phosphosites (Supplementary Table 1). The distribution of modified residues is similar to previous phosphoproteomics studies with 2722 (84%) phospho-serines, 412 (13%) phospho-threonine and 91 (3%) phospho-tyrosines (Figure 1A).

**Figure 1.**
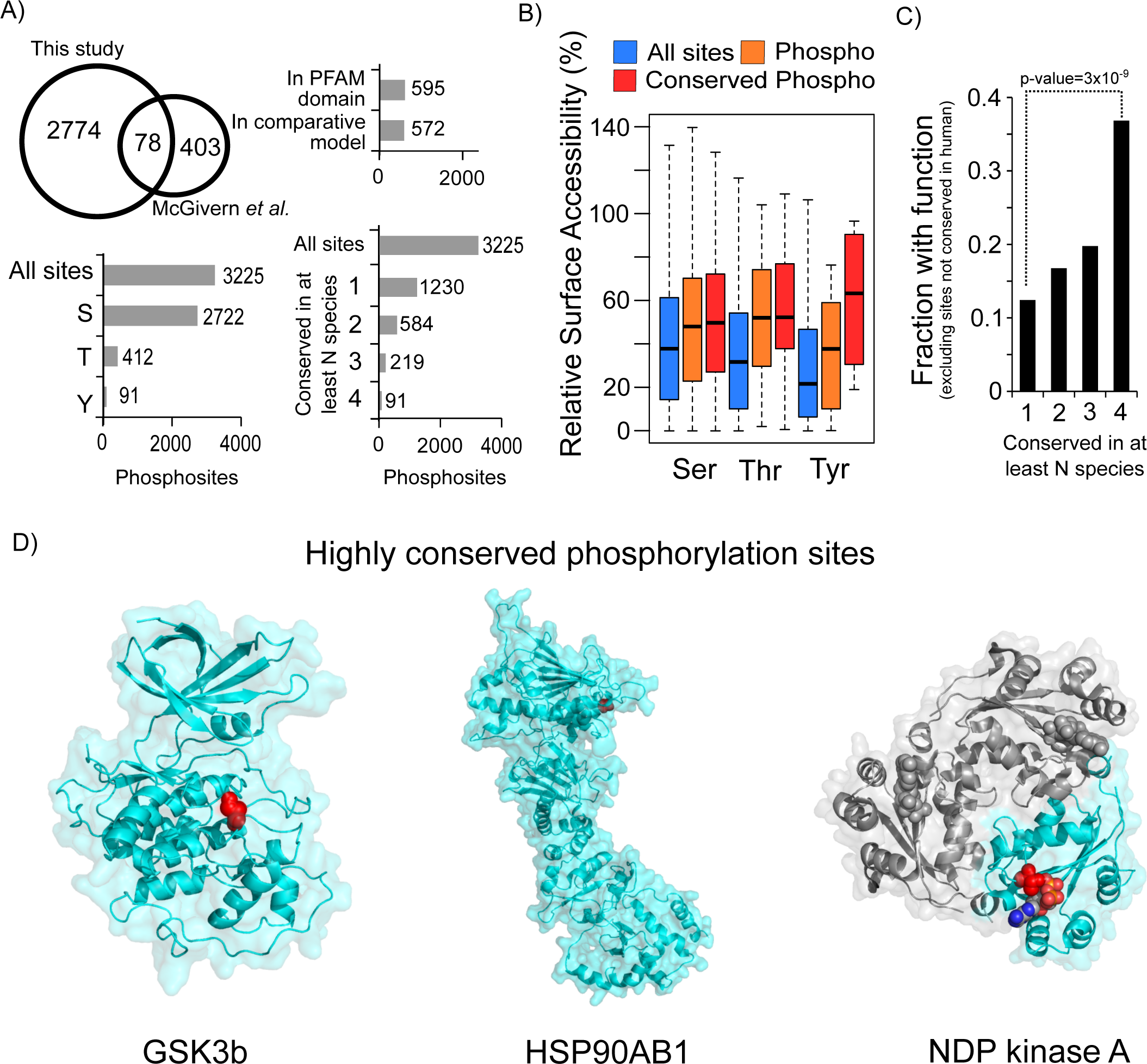
Structural and evolutionary analysis of *X. laevis* phosphosites. A) A total of 3225 non-redundant phosphorylation sites were compiled from the 2852 sites determined here and 481 collected from a previous study [26]. We determined the conservation of these 3225 phosphorylation sites across the 13 other species and obtained structural models for 572 of these sites. B) The all-atom residue relative surface accessibility was compared for all phospho-acceptor residues, phosphosites not conserved or conserved in at least one other species with available phosphorylation data. C) The fraction of *X. laevis* sites with a known function in human increases with the degree of conservation. *X. laevis* sites not conserved in human were excluded from this analysis. D) Example comparative models with highly conserved phosphorylation sites. The phosphorylation site is highlighted in red. For the NDP kinase A, the structure represents the homo-oligomeric complex. One of the subunits is indicated in blue, with the phosphosite position in red and the substrate in the ball-and-stick representation.

In order to study the structural properties of these phosphosites we obtained comparative models for *X. laevis* phosphoproteins. The models were created with ModPipe [27] using templates with at least 25% sequence identity and using established model quality criteria (see Materials and Methods). When different models were available for the same phosphosite-containing region we selected the largest model available. We were able to obtain models for 1239 phosphoproteins covering an average of 46% of the protein length and containing a total of 572 phosphosites. Given that only 595 sites occur within known protein domains as defined in PFAM we believe we are getting structural coverage for a large fraction of the sites found within structured regions of the proteome. In addition to structural information we also determined the level of the conservation of the identified phosphosites using a compilation of phosphorylation information for a set of 13 other species (from ptmfunc.com). *X. laevis* proteins were aligned with putative orthologs in these species and a phosphosite was considered to be conserved in a target species when the aligned peptide region was known to be phosphorylated in that species (see Materials and Methods). Previous studies have noted that regulation by protein phosphorylation can diverge quickly during evolution [4–6,8,9]. In line with these studies we find that only 1230 (38%) sites were found to be conserved in one or more species analyzed (Figure 1A). We note that the conservation values are under-estimated due to lack of complete coverage for most phosphoproteomes. We next combined the structural information and the conservation information to study the surface accessibility of the phosphosites. As expected [28,29], phosphosites are on average more likely to have higher all-atom relative surface accessibility than non-modified serine, threonine and tyrosine residues (Figure 1B). Phosphosites conserved in at least one species don’t appear to be more surface-exposed than average phosphosites (Figure 1B, Serine and Threonine). Conserved phosphotyrosines appear to be more surface-exposed than average phosphotyrosines (Figure 1B) but this difference it is not significant (p-value=0.3, Kolmogorov-Smirnov test) and likely due to a small sample size as only 34 phosphotyrosines are modeled in structures.

A small fraction of phosphosites was found to be conserved across several species. We observed for example that 91 sites were conserved in 4 or more species (Figure 1A and Table 1). In order to test the usefulness of this comparative approach we used a list of human sites known to have a molecular function from small-scale studies (from phosphosite.org). We first restricted the analysis to 700 *X. laevis* phosphosites conserved in *H. sapiens*. We then tested if the level of conservation in additional species beyond human was predictive of a known function in human. While 87 of these 700 (12.4%) *X. laevis* sites have a known human function we observed that the fraction of sites with known function increased with the level of conservation (Figure 1C). Of the 76 Xenopus sites that are conserved in human and in at least 3 other species 28 have a known human function (37%). This large and significant increase suggests that phosphosites conserved across many distantly related species are more likely to be functionally relevant. Increasing coverage of experimentally determined phosphorylation sites across a varied number of species will further facilitate the identification of such highly conserved sites. A list of the *X. laevis* sites that are phosphorylated in the predicted human ortholog and in at least 4 other species are listed in table 1, along with annotations on the molecular role from PhosphositePlus (www.phosphosite.org). Examples of highly conserved sites with available homology models are shown in figure 1C. For example, the phosphorylation of the activation loop region of the GSK3b protein kinase (Figure 1C, left) is one of the most conserved phosphorylation sites across all species. Phosphorylation of the activation loop region of protein kinases is a very well established mechanism to regulate kinase activity. HSP90 proteins are one of the most conserved molecular chaperones involved in the folding of a varied set of client proteins [30]. This chaperone is a highly flexible protein typical forming a homo-dimmer via a c-terminal region [30]. One of the most conserved phosphorylation sites we identified here is in a protein homologous to the human HSP90AB1 (Figure 1C, center). The phosphorylation occurs in the flexible n-terminal region that opens and closes during the ATPase cycle [30]. This position is equivalent to S226 in the human HSP90AB1 that has been previously shown to regulate charperone activity [31]. Another interesting phosphosite, conserved in 4 other species, is the threonine near the active site of NDP kinase A (Figure 1C, right) which due to the proximity to the substrate is very likely to influence enzyme activity. Given the broad phylogenetic distribution of the species used in this study, these highly conserved sites are expected to be enriched in ancient and functionally important phospho-regulatory modifications.

**Table 1.** Highly conserved phosphorylation sites. *X. laevis* phosphosites conserved in human and at least 4 other species are listed with the corresponding human gene name and phosphosite molecular function, if defined in the Phosphosite plus database.

| Protein ID | Phosphosite position(s) | Conserved in N species | Human homolog gene name | Molecular function from Phosphosite plus |
| --- | --- | --- | --- | --- |
| XENLA_tissue_00223572 | 219 | 11 | GSK3B |  |
| XENLA_tissue_00048253 | 219 | 11 | GSK3B |  |
| XENLA_tissue_00292621 | 176 | 8 | CDK7 | Induced kinase activity |
| XENLA_tissue_00119160 | 210 | 8 | HSP90AB1 | Induced HSP90 activity |
| XENLA_tissue_00213365 | 38 | 8 | RPL12 |  |
| XENLA_tissue_00272183 | 304 | 7 | RPLP0 |  |
| XENLA_tissue_00242882 | 438 | 7 | RPLP0 |  |
| XENLA_tissue_00077149 | 226 | 7 | HSP90AB1 | Induced HSP90 activity |
| XENLA_tissue_00277348 | 108 | 7 | EEF1B2 |  |
| XENLA_tissue_00152649 | 245, 246, 250 | 6 | RPS6 | Regulation of molecular association |
| XENLA_tissue_00214244 | 279, 280, 284 | 6 | RPS6 | Regulation of molecular association |
| XENLA_tissue_00157535 | 251 | 6 | CCT3 |  |
| XENLA_tissue_00007232 | 13 | 6 | MCM2 | Induced activity |
| XENLA_tissue_00131108 | 646, 657 | 5 | CLASP1 |  |
| XENLA_tissue_00152649 | 251, 252 | 5 | RPS6 |  |
| XENLA_tissue_00277462 | 14 | 5 | SEC61B |  |
| XENLA_tissue_00237045 | 899, 900, 901 | 5 | GBF1 |  |
| XENLA_tissue_00232516 | 101 | 5 | PRKAR2A |  |
| XENLA_tissue_00172332 | 25 | 5 | MCM2 | Induced activity |
| XENLA_tissue_00007232 | 25,26 | 5 | MCM2 | Induced activity |
| XENLA_tissue_00154464 | 430 | 5 | RPS6KB1 | Induced enzymatic activity |

### Phosphosites are associated with regions with high conformational flexibility

Although phosphosites tend to have high all-atom relative surface accessibility (RSA) around 20% of the sites appear to be poorly accessible - here defined as having below 20% RSA. Given that surface accessibility should be a requirement for kinase regulation we explored potential explanations for these low accessibility sites. We hypothesized that this observation could be due to three potential factors: incorrect homology models, false positive phosphosites, or changes in protein conformation. We reasoned that if model quality was a determinant factor in explaining the inaccessible sites then the fraction of such sites should decrease with increasing quality of the models. However, homology models obtained from templates of higher sequence identify had a similar distribution of phosphosite RSA (Figure 2A). In order to test the impact of false positive sites we relied on the idea that conserved phosphosites are very unlikely to be experimental false positives. We noted that conserved sites have a similar distribution of phosphosite RSA (Figure 2B) than non-conserved sites and only sites that are conserved in 3 or more species have a lower fraction of poorly accessible sites when compared to non-conserved sites (11% vs. approximately 20%, hypergeometric distribution p-value=0.025). Overall these results suggest that false positive phosphosites are unlikely to be a major determinant for low surface accessibility sites.

**Figure 2.**
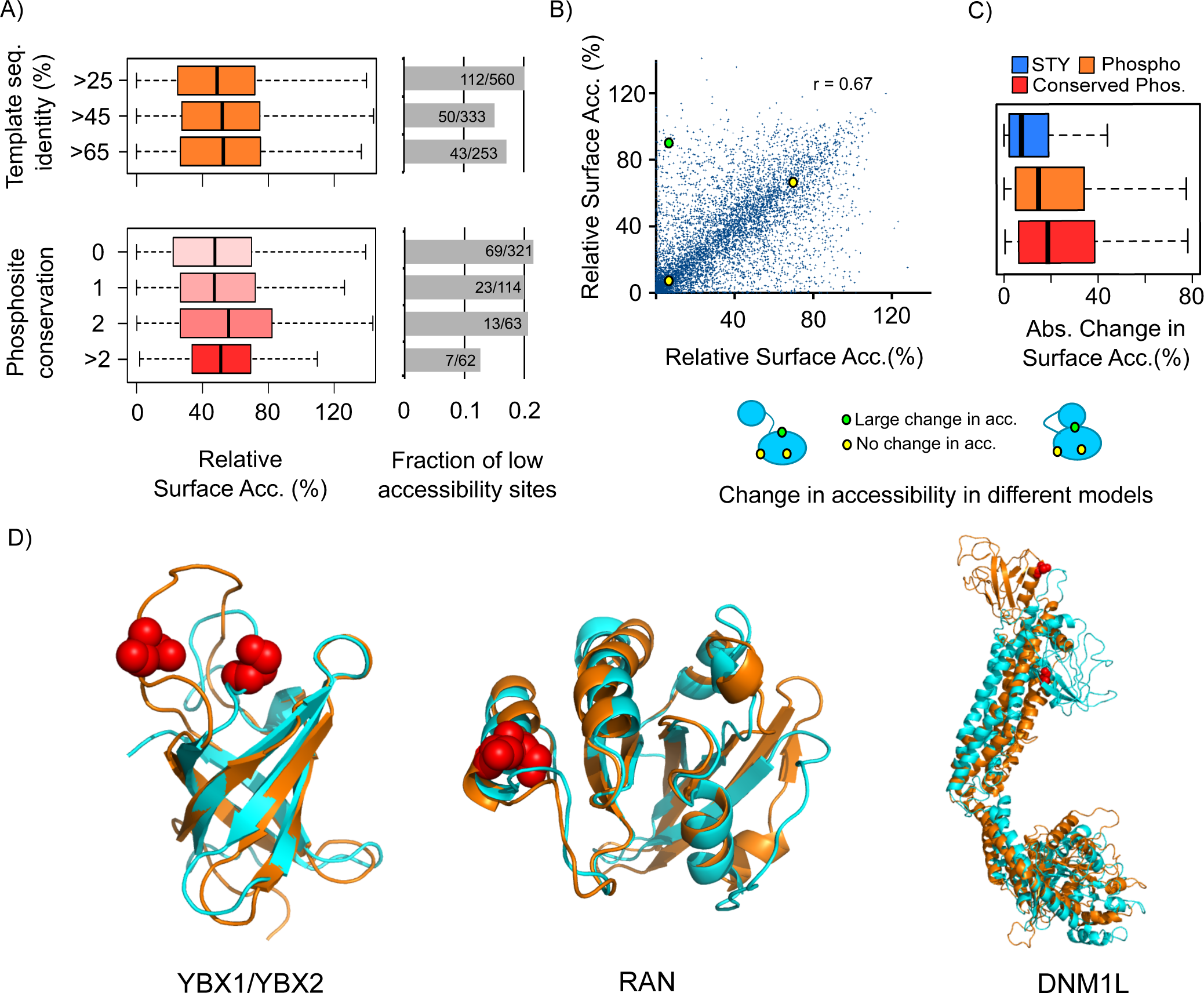
Phosphosites in solvent inaccessible positions may be predictive of conformational flexibility. A) A small fraction of phosphosites (approximately 20%) was observed to be at solvent inaccessible positions (defined here as <20% all-atom RSA). The distribution of phosphosite RSA and the fraction of low accessibility sites do not vary significantly as a function of target-templates sequence identity nor phosphosite conservation. B) For phospho-acceptor residues (Serine, Threonine and Tyrosine) modeled independently based on more than one template structure, we compared the RSA values obtained from different models. These values are highly correlated, although some sites showed large variability in predicted accessibility, potentially indicating regions of conformational flexibility. C) We compared the changes in RSA in different models for phospho-acceptor residues, phosphoryation sites not known to be conserved and those conserved in at least 1 other species. D) Examples of phosphosites found in positions that show a large change in accessibility in two templates and are poorly accessibility in one of the templates. The phosphosite position is highlighted in red and the models in orange have a higher RSA for the phosphosite position when compared to the model in cyan.

If inaccessible phosphosites are not mostly due to incorrect homology models or false positive sites then conformation flexibility might explain why some sites appear to have low accessibility. The regions that are modified might exist in different conformations and the phosphorylation occurs in a more accessible conformation that is not captured in the structure template used to create these models. In order to study this we analyzed phosphoproteins for which we had more than one model structure obtained from different templates. For each pair of comparative models we correlated the accessibility for all serine, threonine and tyrosine residues. Overall, there is a high correlation of all-atom RSA value for different templates (Figure 3C, r=0.67). However, phosphorylated residues showed a significantly higher change in accessibility in different models when compared to non-modified residues (Figure 3C, p-value=4x10^−4^, Kolmogorov-Smirnov test). The median absolute change in accessibility is 7.4 for the phospho-acceptor residues, 13.5 for phosphosites and 18.35 for conserved phosphosites. This result suggest that phosphosites are more likely to be in regions that show high conformation flexibility. Phosphorylation can, in some cases, regulate protein conformation [32] and this analysis suggests that it is possible to use comparative models to define a class of functional phosphosites that can play a role in conformation regulation.

**Figure 3.**
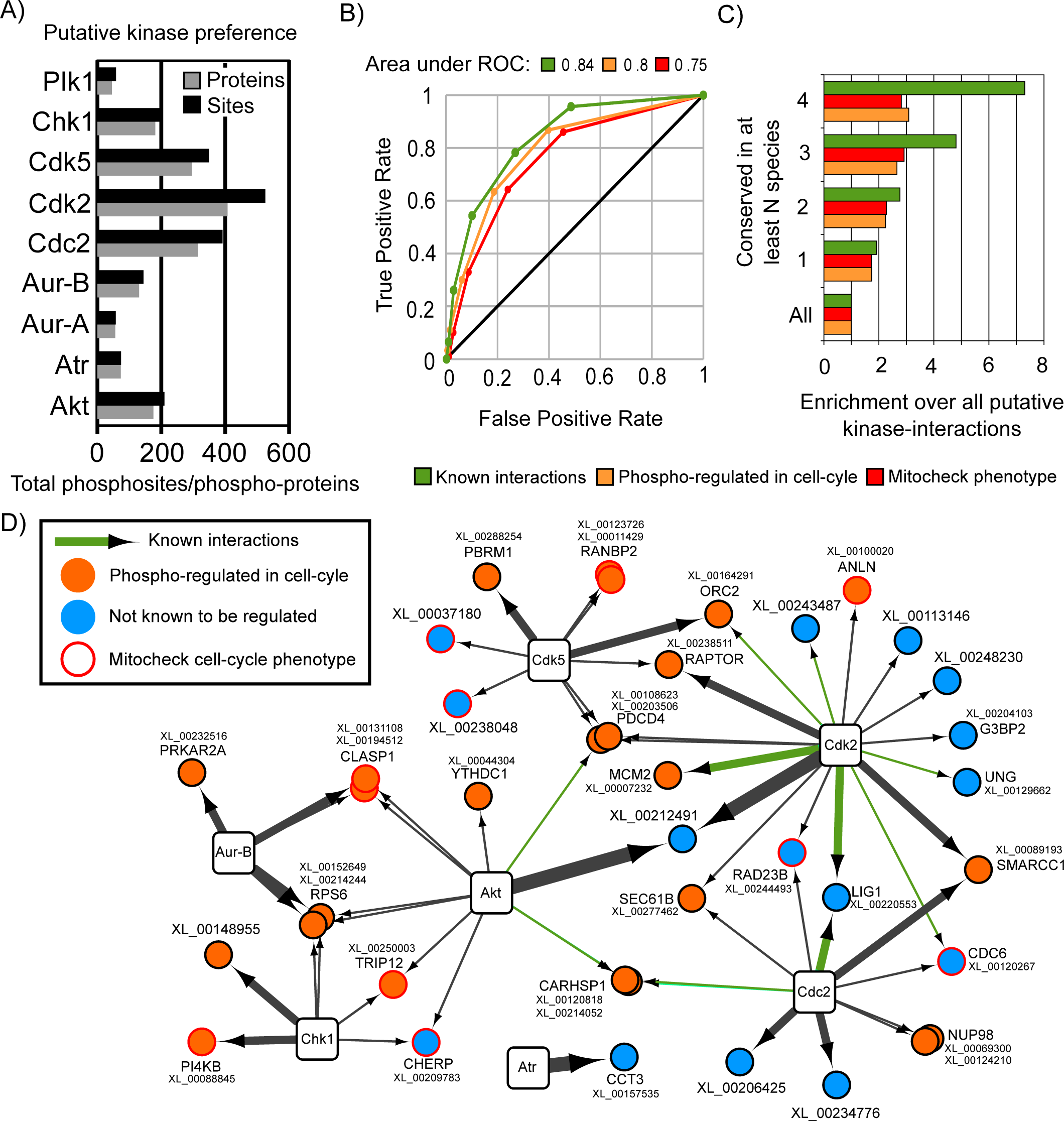
Conservation of putative cell-cycle related kinase interactions is predictive of know and/or cell-cycle related kinase-target interactions. A) The number of predicted kinase target sites and proteins associated with 1 of 9 cell-cycle kinases selected for analysis in *X. laevis*. We tested if the degree of conservation of kinase-interactions was predictive of known interactions; enriched in proteins that are phospho-regulated in the cell cycle; and genes known to cause cell cycle phenotypes when knocked down. B) ROC curves measuring the accuracy for kinase-interaction predictions. C) Enrichment over random prediction for the 3 tested features. D) Predicted kinase interactions conserved in 4 or more species are shown, highlighting known interactions, proteins phospho-regulated during the cell cycle and genes causing cell cycle phonotypes.

We analyzed in more detail sites that were at positions with large changes in accessibility across different templates and where the site had RSA below 20% in at least one model. Three of such cases are shown in Figure 3E were we superimposed the two models highlighting the differences in accessibility. YBX1 and YBX2 are RNA binding proteins and both are phosphorylated in the cold-shock protein (CDP) domain (PFAM:PF00313) in a loop region that is known to be highly flexible [33]. This domain can bind directly RNA and we speculate that the addition of a negative charge in this loop could influence RNA binding. The Ran GTPase was also observed to be phosphorylated in a position with different accessibility in different templates (Figure 2D, center). The phosphosite is in an position equivalent to S135 in human RAN that has been previously shown to be regulated by p21-activated kinase 4 (PAK4) during the cell-cycle in human and *X. laevis* [34]. In addition, Ran S135 alanine and phosphomimetic mutants had an impact on microtubule nucleation in *X. laevis* extracts and in binding the exchange factor RCC1 in human cell lines [34]. It is possible that PAK4 regulation of S135 could promote a conformation that does not favor the interaction with RCC1. The third example is a phosphosite on the Dynamin 1-Like pleckstrin homology (PH) domain that comprises the “foot” region of dynamin like proteins (Figure 3D, right). The PH domain can adopt different conformations relative to the rest of the protein and the phosphorylated position changes drastically in accessibility depending on the conformation. The phosphorylated position is equivalent to the S635 in rat Drp1 and S616 in human Drp1 that has been previously shown to be regulated during cell-cycle and have functional roles in mitochondrial fission and microtubule targeting [35,36]. We hypothesize that the phosphosite may regulate Drp1 function by restricting the possible conformation variability of the PH domain relative to the rest of the protein. Although these examples would require experimental validation they illustrate how the structural analysis of phosphosites using different structural templates might suggest mechanist explanations for the function of MS identified sites.

### Degree of conservation in predicted kinase-protein interactions correlates with known and functionally meaningful phospho-regulatory events

Previous studies have shown that the conservation of kinase motifs in alignments of orthologous proteins could be used to improve the predictions of kinase target sites [15,16]. However, some kinases are known to regulate proteins in clusters of sites [37–39] and in some cases it is plausible that the exact position of the target site might change within the protein during evolution yet maintaining the kinase-protein regulatory interaction. In line with this reasoning, predicted kinase-protein interactions were often found to be conserved across species even when the phosphosite positions were not conserved [6]. We hypothesize that the conservation of predicted kinase-protein interactions across the large number of species analyzed here could be used to predict functionally important interactions. We focused on a set of 10 cell cycle-related kinases for which specificity models have been described and a list of known kinase-protein interactions was available for benchmarking (see Materials and Methods). Focusing on cell cycle kinases also allowed us to take advantage of previous large scale studies of cell cycle phosphoregulation [40] and phenotypes [41] for benchmarking purposes.

For each kinase, substrate recognition models were obtained from GPS and all phosphorylated residues in all species were scored against these models. Kinase-site scores were Z-score normalized and benchmarked against a set of known human kinase target sites (see Materials and Methods). We used the area under the ROC curve (AROC) as a metric to evaluate the prediction accuracy of these models (Supplementary Table 2). Across all kinases, the AROC value was 0.86 suggesting that these models can successfully discriminate between different kinase binding preferences. We excluded CDK7 from subsequent analysis given that the performance of this kinase model was below average (AROC=0.63). For each species we predicted which phosphosites match the kinase preference of the 9 kinases studied. At the threshold used (z-score>4) we estimate a coverage of 30% and a 6-fold enrichment over random of true positive target sites (see Materials and Methods). A phosphoprotein was predicted to be regulated by a kinase if at least one phosphosite was a predicted target of that kinase. The number of *X. laevis* sites and proteins predicted to be regulated by each kinase is shown in Figure 3A.

We next tested if the degree of conservation of kinase interactions is a useful predictor for known and functionally important kinase-target interactions. A putative *X. laevis* kinase-protein interaction was considered to be conserved in another species if an ortholog in that species had at least one phosphosite that was predicted to be a target of the same kinase model. We note that the sites do not have to be in the same region of the orthologous proteins. We ranked *X. laevis* kinase interactions according to the degree of conservation in the other 13 species. We observed that the degree of conservation was a significant predictor of previously described human kinase-protein interactions (Figure 3B AROC=0.84 and 3C, “known interactions”). Highly conserved kinase-protein interactions were also enriched in target proteins that were previously known to be phosphoregulated during the cell cycle in human [40] (Figure 3B AROC=0.8 and 3C, phospho-regulated) and enriched in genes that when knocked-down cause cell cycle-related phenotypes [41] (Figure 3B AROC=0.75 and 3C, mitocheck phenotype). We compared this approach with a sequence based method. For each species we used the kinase matrices to predict known kinase-protein interactions using also a z-score of 4 but scoring any acceptor residue and disregarding the phosphoproteomic information. When compared to the method described here sequence based conservation alone resulted in a much less significant enrichment in known kinase-protein interactions (AROC=0.69 vs 0.84).

We selected the putative kinase-interactions conserved in 4 or more species for further analysis (Figure 3D). When compared to all putative kinase-interactions this network is 7.3-fold enriched in known kinase-interactions (Figure 3C and 3D green arrows), 3-fold enriched in cell cycle phospho-regulated proteins (Figure 3C and 3D orange circles) and 2.8-fold enriched in genes associated with cell cycle-related phenotypes (Figure 3C and 3D red outline). We also searched the literature for evidence supporting some of the predicted interactions not yet captured in the Phosphosite database. For example, the predicted regulation of the nuclear pore protein Nup98 by Cdc2 is supported by experimental evidence in human cells [42] where Nup98 has been shown to be phosphorylated by Cdc2, among other kinases. In addition, the phosphorylation of this protein has an impact on the disassembly of the nuclear pore [42]. Cdc6 has also been shown to be an in vitro target of Cdc2 in *X. laevis* egg extracts [43] and that CDK activity promotes the release of Cdc6 from chromatin an important mechanism to regulate the preinitiation complex. Although we could not find direct evidence for the regulation of raptor by Cdk2, it has been shown that this protein is phosphorylated in human cells during mitosis in a Cdk1 dependent fashion [44]. This regulation is important for cell cycle progression and regulation of mTOR activity [44]. These examples and the conservation analysis described above strongly suggest that kinase specificity models can be used in comparative phospho-proteomics analysis to improve the prediction of kinase targets. A list of the putative interactions from Figure 3D with conservation information is available in supplementary table 3 to facilitate further studies.

## Discussion

To facilitate the study of *X. laevis* PTM signaling, we obtained here a large-scale survey of this species’ phosphoproteome. Previous studies have indicated that phosphoregulation can diverge quickly during evolution and that a fraction of phosphosites might serve no biological function in extant species. In order to increase the usefulness of the obtained phosphosites we used a compilation of phosphorylation data for 13 other species to identify highly conserved phosphosites and potential kinase-protein interactions. While others have shown that conservation of kinase sequence motifs is a useful filter to predict kinase interactions [15,16] we show here that the degree of conservation of experimentally determined phosphorylation states is a stronger predictor of sites with known function, of kinase-protein interactions and of function specific annotations (i.e. cell-cycle regulated and phenotypes). A small fraction of sites appear to be conserved over a large number of species. Given the divergence times separating the species studied here, these phosphosites likely have very ancient origins despite the fast evolutionary turn-over of phospho-regulation. As the experimental data on protein phosphorylation is incomplete for most species the conservation values presented here are under-estimated. Additional data will allow for further identification of such “ultra-conserved” and functionally important sites. However, we note that although conservation is useful predictor of functional phosphosites, there are functionally important sites that are not highly conserved. For these reasons it is important to develop approaches that do not rely on conservation to rank PTMs according to biological importance. In this context, we and others have previously made use of structural information to predict PTMs that have the potential to regulate protein-protein interactions [10,17]. Using comparative models for *X. laevis* phosphoproteins we observed that some phosphosites appear to be in inaccessible regions and that phosphosites tend to be in positions of higher variability in surface accessibility across different structural templates. Our analysis suggests that structural models can therefore be used to predict, in an unbiased way, PTMs with the potential to regulate protein conformation. This approach could be further extended by computational methods that take into account molecular dynamics [45].

The majority of PTMs identified to date for human and other species has no known function. Given the large throughput of MS approaches and the low fraction of PTMs with currently known functions much additional effort needs to be committed to the development of computational and experimental methods to elucidate PTM function. The evolutionary and structural observations presented here can be used to facilitate the prioritization of PTM functional studies in any species.

## Materials and Methods

### Xenopus Extract Preparation

Interphase egg extracts supplemented with an ATP regenerating system and cycloheximide (100mg/ml) were prepared as described [46,47]. In order to make mitotic egg extracts, interphase egg extracts were treated with 100nM non-degradable Xenopus D65-cyclin B1 and reactions were incubated for 1 hour and 30min at 22C. De-membranated sperm chromatin was added at 500/μl and samples were collected and stained with DAPI in order to monitor nuclear morphology and mitotic progression by fluorescence and phase microscopy. The criteria for M phase entry were condensed chromatin and a lack of a discernable nuclear envelope in at least 90% of the nuclei.

### Sample preparation for mass spectrometry analysis

Xenopus extracts were denatured in a buffer containing 8M urea, 0.1M Tris pH 8.0, and 150 mM NaCl. Disulfide bonds were reduced by incubation with 4 mM TCEP for 30 minutes at room temperature, and free sulfhydryl groups were alkylated by incubation with 10 mM iodoacetamide for 30 minutes at room temperature in the dark. Samples were diluted back to 2 M urea by addition of 0.1 M Tris pH 8.0, and trypsin was added at an enzyme:substrate ratios of 1:100. Lysates were digested overnight at 37 degrees Celsius. Following digestion the samples were concentrated using SepPak C18 cartridges (Waters). The C18 cartridge was washed once with 1 mL of 80% ACN, 0.1% TFA, followed by a 3 mL wash with 0.1% TFA. 10% TFA was added to each samples to a final concentration of 0.1% after which the samples were applied to the cartridge. The cartridge was washed with 3 mL of 0.1% TFA after binding, and the peptides were eluted with 40% ACN, 0.1% TFA. Following elution the peptides were lyophilized to dryness. Phosphopeptides were fractionated using hydrophilic interaction chromatography (HILIC) adapted from a method published by McNulty and Annan [48]. Buffers used for HILIC separation were HILIC buffer A (2% ACN, 0.1% TFA) and HILIC buffer B (98% ACN, 0.1% TFA). Peptides were resuspended in 90% HILIC buffer B and loaded onto a TSKgel amide-80 column (Tosoh Biosciences, 4.6 mm I.D. x 25 cm packed with 5 um particles). Peptides were separated at a flow rate of 0.5 mL / min using a gradient from 90% to 85% HILIC buffer B for 5 minutes, 85% to 55% HILIC buffer B for 80 minutes, then 55% to 0% HILIC buffer B for 5 minutes. Fractions were collected every 2 minutes and the 22 fractions previously determined to contain the majority of phosphopeptides were evaporated to dryness. Following HILIC fractionation, fractions were further enriched for phosphopeptides using titanium dioxide magnetic beads (Pierce) using the manufacturer’s protocol. Following titanium dioxide enrichment, sample were evaporated to dryness and resuspended in 0.1% formic acid for mass spectrometry analysis.

### Mass spectrometry analysis and phosphosite identification

Each fraction was analyzed separately by a Thermo Scientific LTQ Orbitrap Elite mass spectrometry system equipped with an Easy-nLC 1000 HPLC and autosampler. Samples were injected directly onto a reverse phase column (25 cm x 75 um I.D. packed with ReproSil-Pur C18-AQ 1.9 um particles) in buffer A (0.1% formic acid) at a maximum pressure of 800 bar. Peptides were separated with a gradient from 0% to 5% buffer B (100% ACN, 0.1% formic acid) over 5 minutes, then 5% to 30% buffer B over 52 minutes, then 30% to 95% buffer B over 1 minute, then held at 95% buffer B for 6 minutes. The separation was performed at a flow rate of 400 nl/min. The mass spectrometer continuously collected spectra in a data-dependent manner, acquiring a full scan in the Orbitrap (at 120,000 resolution with an automatic gain control target of 1,000,000 and a maximum injection time of 100 ms) followed by collision-induced dissociation spectra for the 20 most abundant ions in the ion trap (with an automatic gain control target of 10,000, a maximum injection time of 10 ms, a normalized collision energy of 35.0, activation Q of 0.250, isolation width of 2.0 m/z, and an activation time of 10.0). Singly and unassigned charge states were rejected for data-dependent selection. Dynamic exclusion was enabled to data-dependent selection of ions with a repeat count of 1, a repeat duration of 20.0 s, an exclusion duration of 20.0 s, an exclusion list size of 500, and exclusion mass width of + or −10.00 ppm. Raw mass spectrometry data was converted to peaklists using the PAVA algorithm. Data were searched using the Protein Prospector suite of algorithms (prospector.ucsf.edu). The data were searched against a X. laevis proteome obtained from the genome sequencing project. Specifically, we used a version containing 24,762 gene models obtained from sequencing of tissue samples and containing the longest gene model for each putative orthologous group. ("OrthoGeneOne" model of Taira201203_XENLA_tissue data, available at http://www.marcottelab.org/index.php/XENLA_GeneModel2012). Searches were run with a concatenated decoy database comprised of all sequences with their amino acids randomized. The algorithm searched for fully tryptic peptides with up to 2 missed cleavages using a parent mass tolerance of 20 ppm and a fragment mass tolerance of 0.8 Da. The algorithm indicated a static modification for carboxyamidomethyl of cysteine residues, and for variable modifications acetylation of protein N-termini, glutamine to pyroglutamate conversion, methionine oxidation, and phosphorylation of serine, threonine, or tyrosine residues. Data were filtered using a Protein Prospector expectation value that was resulted in a false positive rate of 1% as determined by the number of matches to the randomized decoy database. The list of identified phosphopetides are provided in Supplementary Table 1.

### Data analysis

Putative orthologs of *X. laevis* phosphoproteins were predicted using the reciprocal best-BLAST hits method [49] against a set of 13 proteomes with currently available phosphorylation information. Putative orthologs were aligned using MUSCLE [50]. The phosphorylation information for the 13 species was retrieved from the ptmfunc database (http://ptmfunc.com) and includes phosphosite information for *Saccharomyces cerevisiae*, *Schizosaccharomyces pombe*, *Plasmodium falciparum*, *Toxoplasma gondii*, *Trypanosoma brucei*, *Trypanosoma cruzi*, *Oryza sativa*, A*rabidopsis thaliana*, *Drosophila melanogaster*, *Caenorhabditis elegans*, *Homo sapiens*, *Rattus norvegicus* and *Mus musculus.* An *X. laevis* phosphosite was considered to be conserved in a target species if the predicted orthologous protein was known to be phosphorylated in the target species in a window of +/-2 residues around the aligned position. A window was used to take into account the alignment uncertainty and the ambiguity in identifying the exact position of the phosphorylated residue within a phosphopeptide. To predict the targets of cell cycle-related kinases, we selected 10 kinases (Akt, Atr, Chk1, Cdc2, Cdk2, Cdk5, Cdk7, Aur-A, Aur-B and Plk1) with kinase substrate models available in GPS and with more than 30 known human phosphosite targets (obtained from the PhosphositePlus database) for benchmarking purposes. For each kinase, we tested the ability of the GPS kinase model to correctly discriminate between the known target sites and the target sites of the other 9 selected kinases. The area under the ROC curve was used as the accuracy criterion. With the exception of Cdk7, all kinases had an AROC value greater than 0.74; thus, we excluded the Cdk7 model from further analysis. The phosphosites of all 14 species were scored against the GPS kinase models. The GPS model output values were normalized to obtain z-scores for each kinase; a Z-score cut-off of 4 was used to predict the kinase-site interactions. Using the benchmark set of known target sites, we estimate that this cut-off leads to approximately 30% coverage of and 6-fold enrichment of true positives over a random selection. Structural models of *X. laevis* phosphoproteins were built automatically using ModPipe [27] relying on Modeller 9.10 [51]. A model was considered acceptable if the template sequence identify was at least 25% and met one additional criterion: TSVMod NO35 >=40%, GA341 >=0.7, E-value <0.0001 or zDOPE <0. All-atom residue relative surface accessibility was computed using NACCESS [52].

## Acknowledgments

S.D.M.S. is supported by an MRC career development award. P.B. is supported by the Human Frontier Science Program (CDA00069/2013-C). A.S. and U.P. are supported by the NIH NIGMS grant U54 GM074945. We thank the Xenopus laevis genome project consortium to provide gene annotation information from unpublished RNA-seq data. Especially, for the RNA-seq based gene model we used in this project, we thank Shuji Takahashi, Atsushi Toyoda, Yutaka Suzuki, Sumio Sugano, Asao Fujiyama, and Masanori Taira for sharing their unpublished RNA-seq data (the construction of RNA-seq data sets was supported in part by KAKENHI (Grant-in-Aid for Scientific Research) on Innovative Areas "Genome Science" from the Ministry of Education, Culture, Sports, Science and Technology of Japan), and Taejoon Kwon, Shuji Takahashi, Toshiaki Tanaka, Edward Marcotte for gene model construction and validation.

## Supporting Information Legends

Supplementary table 1 – List of experimentally identified phospho-peptides. Mass-spectrometry derived phosphopeptides are described in accompanying spreadsheet.

Supplementary table 2 – Kinase models tested. 10 kinases were selected for predictions based on their functional associations with the cell cycle, the availability of a kinase specificity model in the Group Based Prediction system (http://gps.biocuckoo.org/) and at least 30 known kinase-site interactions described in the PhosphositePlus database (http://phosphosite.org). For each kinase we tested the capacity of the model to descriminate between the known targets and other phosphosites known to be regulated by protein kinases. AROC – area under the receiver operating characteristic curve. Pos – number of positive kinase-site interactions scored. Neg – Number of negative kinase-site interactions scored. The model for CDK7 was excluded from further analysis given its relatively low prediction accuracy.

Supplementary table 3 – List of predicted kinase interactions conserved in 4 or more species. Cell cycle regulated: human protein known to have phosphosites that are regulated during the cycle (1 – yes; 0 – no). Mitocheck pheno:human gene known to cause a mitotic phenotype as described in the Mitocheck database (1 – yes; 0 – no); TP-Known human kinase-substrate interaction as described in the PhosphositePlus database (1 – yes; 0 – no).Conservation: Number of species with a conserved predicted kinase-protein interaction.

